# HARIBOSS: a curated database of RNA-small molecules structures to aid rational drug design

**DOI:** 10.1101/2022.05.17.492306

**Authors:** F. P. Panei, R. Torchet, H. Menager, P. Gkeka, M. Bonomi

## Abstract

RNA molecules are implicated in numerous fundamental biological processes and many human pathologies, such as cancer, neurodegenerative disorders, muscular diseases, and bacterial infections. Modulating the mode of action of disease-implicated RNA molecules can lead to the discovery of new therapeutical agents and even address pathologies linked to 8undruggable9 protein targets. This modulation can be achieved by direct targeting of RNA with small molecules. As of today, only a few RNA-targeting small molecules are used clinically. One of the main obstacles that has hampered the development of a rational drug design protocol to target RNA with small molecules is the lack of a comprehensive understanding of the molecular mechanisms at the basis of RNA-small molecule recognition. Here, we present HARIBOSS, a curated collection of RNA-small molecule structures determined by X-ray crystallography, Nuclear Magnetic Resonance spectroscopy and cryo-electron microscopy. HARIBOSS facilitates the exploration of drug-like compounds known to bind RNA, the analysis of ligands and pockets properties, and ultimately the development of *in silico* strategies to identify RNA-targeting small molecules. HARIBOSS can be explored via a web interface available at http://hariboss.pasteur.cloud.

## Introduction

During the past two decades, RNA molecules have been shown to perform a variety of vital biological functions besides being a passive carrier of genetic information from DNA to protein. An explosion of research in the field of RNA biology has provided information about RNA diversity with several new definitions of RNA types as well as structural and functional information. For example, it is now well established that RNA is implicated in the regulation of gene expression at the levels of transcription, RNA processing and translation, in the regulation of epigenetic modifications and in the protection of the nucleus from foreign nucleic acids [1-4]. In conjunction with these discoveries, modulating RNA function is emerging as a promising therapeutic approach against pathologies such as cancer, viral infections, cardiovascular and muscular diseases, and neurodegenerative disorders [5, 6]. Traditionally, RNA modulation has been achieved using oligonucleotides such as small interfering RNA (siRNA), antisense, aptamers, and other RNA moieties or direct RNA-editing by CRISPR-Cas9 [7-10]. While oligonucleotides have been successful in binding to and modulating RNA, their drug bioavailability and membrane penetration have been quite challenging. Moreover, part of these molecules carry a large negative charge and, therefore, are susceptible to degradation by RNAses [11, 12]. Small molecules able to selectively bind to RNA provide a more attractive alternative from a bioavailability and delivery perspective [13-17].

Direct RNA targeting with small molecules has the potential to be revolutionary. Only 1.5% of the human genome encodes proteins and only a small fraction (12%) of this percentage is related to diseases and targeted by existing drugs (3%). Strategies like targeting non-coding RNAs, which represent instead the majority of the human genome, or the mRNA of undruggable proteins will therefore allow to significantly expand the space of potential targets [18]. Several small molecules that interfere with RNA functions have already been identified [19-23], suggesting that therapeutics based on small-molecules targeting RNA may be possible. However, only a few compounds have been approved so far by the United States Food and Drug Administration (FDA), namely linezolid, ribocil and risdiplam. Linezolid (Zyvox), initially discovered in the mid 1990s and approved for commercial use in 2000, is a broad-spectrum antibacterial agent that binds to the large RNA subunit of the ribosome and interferes with the positioning of the tRNA [24]. Ribocil, discovered by Merck in 2015, is a drug that selectively binds the flavin mononucleotide (FMN) riboswitch (RNA-mediated regulator of gene expression in bacteria) and silences gene expression, making it effective in the treatment of bacterial infections [25]. Interestingly, ribocil binds in the same pocket as the natural flavin mononucleotide ligand. Risdiplam, discovered by Roche-Genentech in 2018, is a drug that modulates the splicing of the Survival Motor Neuron 2 (SMN2) mRNA and mitigates the pathological SMN2 protein states related to Spinal Muscular Atrophy [26]. All these three compounds as well as most of those under pre-clinical or clinical evaluation have been discovered using loss/gain-of-function studies, phenotypic screening, or animal models.

Computer-aided drug design (CADD) has the potential to guide the rational development of small molecules targeting RNA [27, 28]. To date, this strategy is hampered by our limited understanding of RNA structural and dynamic properties as well as of the mechanisms of RNA-small molecule (RNA-SM) recognition [14, 18]. Previous efforts to characterize the physico-chemical properties of RNA binders and their intersection with the chemical space of drugs targeting proteins provided insights into their molecular interaction with RNA [16, 29-31]. Databases that collect all the known compounds binding RNA can be exploited for ligand- [32, 33] and 2D structure-based [34-36] virtual screening. DrugPred_RNA is, to the best of our knowledge, the only tool that has been trained and tuned using 3D structure data in order to characterize RNA binding sites [37]. In addition, most of current CADD pipelines have been developed for protein targets and might not be directly applicable to RNA. A first step in the development of structure-based approaches therefore requires an extensive assessment of the existing tools for pocket detection and ligand docking and possibly the development of new tools that exploit all the available structural information on RNA-SM complexes. While there have been previous efforts to collect such data [38-41], there is currently no comprehensive, curated, and regularly updated repository available to the scientific community.

Here, we present “Harnessing RIBOnucleic acid 3 Small molecule Structures” (HARIBOSS), a curated online database of RNA-SM structures. Each entry in HARIBOSS corresponds to a structure deposited in the RCSB Protein Data Bank (PDB) database [42] containing at least one chemical compound matching a list of basic drug-like criteria and interacting with at least one RNA chain. Ligands are annotated by their physico-chemical properties and by the number and composition of interacting RNA molecules. RNA pockets occupied by ligands are characterized in terms of geometric properties, such as volume, hydrophobicity and hydrophilicity, and overall propensity to bind small molecules and drug-like compounds. HARIBOSS is freely accessible via a web interface (http://hariboss.pasteur.cloud) and will facilitate understanding the nature of the interactions that drive RNA-SM recognition and benchmarking existing tools for *in silico* drug design with RNA targets.

## Materials and Methods

### Construction of the database

The HARIBOSS database was built in 3 steps (**Fig. 1A**), which were implemented in a series of python scripts. The operations described below are performed on a monthly basis to update HARIBOSS with the new structures deposited in the PDB.

**Figure 1.**
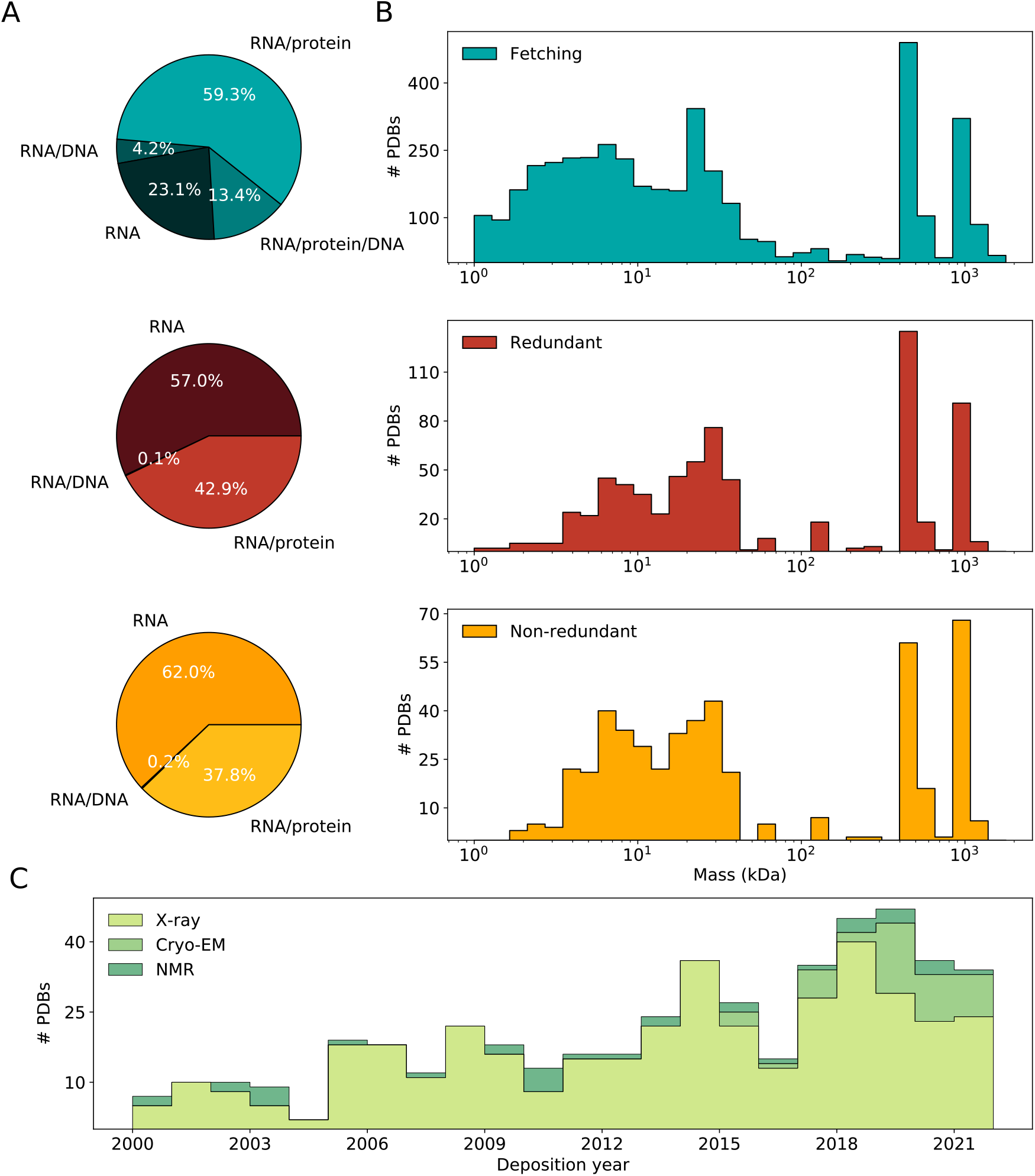
General properties of the RNA-SM structures included in the HARIBOSS database. A) Composition of HARIBOSS in terms of RNA, RNA-protein, RNA-DNA and RNA-DNA-protein structures at the different steps of the database construction: initial fetching from the PDB database (cyan), filtering based on ligand composition and nature of the interacting partners (redundant HARIBOSS, red), clustering based on sequence and pocket structure similarity (non-redundant HARIBOSS, yellow). B) Distribution of the total mass of the system at the three different steps of HARIBOSS construction. C) Number of PDBs in the non-redundant HARIBOSS resolved by X-ray crystallography (X-ray, light green), cryo-electron microscopy (cryo-EM, green) and Nuclear Magnetic Resonance spectroscopy (NMR, dark green) as a function of the deposition year.

#### I. Initial fetching from the PDB

The first step consisted of querying the PDB and collecting all the structures that contained at least one RNA molecule and one ligand. This operation was performed using a solr-based search API developed by the European Molecular Biology Laboratory of the European Bioinformatics Institut (EMBL-EBI) [43]. At this stage, we collected structures in which small molecules interact with RNA, DNA, proteins or a combination of these biological entities.

#### II. Identifying structures with drug-like compounds bound to RNA

Among the initially fetched structures, we selected those that contained drug-like compounds non-covalently bound only to RNA. To accomplish this, we adopted the following procedure. For each entry, the correspondent mmCIF file was processed by MDAnalysis v. 2.0.0-beta [44] and OpenMM v 7.5.1 [45] to classify its constituents into RNA, DNA, protein, ions, and water molecules. Modified residues were assigned to RNA, DNA, or protein molecules based on the information about covalent bonds retrieved from the mmCIF with Biopython v. 1.79 [46]. All the molecules not included in the categories defined above were classified as ligands. We considered a ligand to be a drug-like compound if it satisfied the following criteria [38]:

- mass within 160 and 1000 Da, as reported in the mmCIF file [46];
- presence of at least one C atom;
- presence of only C, H, N, O, Br, Cl, F, P, Si, B, Se atoms;

For each compound fulfilling these criteria, we defined as interacting all the molecules within 6 Å from the ligand atoms. RNA chains with less than 10 atoms in the radius of 6 Å around the ligand were not considered as interacting (**Table S1**). For the first release of HARIBOSS, we retained in the database only the structures in which ligands interact exclusively with RNA chains. At this stage, we annotated each entry with the following information: total mass of the system, molecular composition (RNA, RNA/DNA, RNA/protein, RNA/DNA/protein), experimental method used to determine the structure (experiment type, resolution and deposition year), properties of bound ligands (PDB name, residue number and chain id, molecular mass, identity of the interacting RNA chains). The RNA-SM complexes obtained at the end of this filtering step constitute the *redundant* version of the HARIBOSS database.

#### III. Clustering of RNA/ligand complexes

We defined a clustering procedure to select representative RNA/ligand complexes and build a *non-redundant* version of HARIBOSS. Since the database contained ligands interacting with more than one RNA chain, we adopted the following procedure:

1. we created a FASTA file with all the sequences of the individual RNA chains, each annotated with its interacting ligand, which we processed with CD-HIT-EST [47] to cluster them at 90% sequence identity;
2. in case of ligands interacting with multiple RNA chains, we defined two structures to belong to the same cluster if the individual chains from the two entries were clustered together at step 1;
3. to obtain a variety of different RNA/ligand interactions, RNA chains belonging to the same cluster but bound to different ligands were classified in separate subclusters;
4. these subclusters were further classified based on the structural similarity of the pocket atoms by performing a hierarchical clustering using the RMSD of the aligned pocket atoms of the nucleic backbone as metrics and a cutoff of 1.5 Å;
5. for each cluster, we selected as representative the entry closest to the cluster center with highest experimental resolution, which we consider an appropriate choice for the use of HARIBOSS in computational structural studies.

### Analysis of the database

The HARIBOSS entries were analyzed based on general information about the structure, physico-chemical properties of the ligands and of the RNA pockets.

#### Exploration of the chemical space of RNA binders

To present an overview of the chemical space of the RNA binders included in our database, the TMAP visualization method was used [48]. In TMAP the molecules are represented by their fingerprints and indexed in an LSH forest structure. Based on the distances calculated during this step, an undirected weighted *c*-approximate *k*-nearest neighbor graph (*c*-*k*-NNG) is used to construct a minimum spanning tree. This tree is then projected onto the Euclidean plane. For the creation of the spanning tree, the MHFP6 fingerprints and a point size of 20 were used. For more information about the method, we refer the readers to the original publication [48]. TMAP was chosen over simple clustering or other chemical space visualization tools for the informative nature of its tree-based layout, which enables to locally as well as globally locate specific clusters and their relative position compared to other clusters. Moreover, such an approach will be able to accommodate in the future the expanding chemical space of RNA binders.

#### Analysis of ligand properties

QikProp (Schrodinger Suite 2021.v3) was employed for the calculation of the following ligand properties: solvent-accessible surface area (SASA) and its hydrophobic and hydrophilic content (FOSA and FISA respectively), predicted IC50 value for blockage of HERG K+ channels, Caco-2 cell permeability and brain/blood partition coefficient. Furthermore, the volume of each ligand was calculated using the calc_volume.py tool included in Schrodinger Suite 2021.v3.

#### Cavity volume analysis with mkgridXf

To identify the cavities occupied by ligands in the structures of our HARIBOSS database and measure their volume, we followed a 3-steps procedure.

1. Cavity identification. We started by using mkgridXf [49] to identify potential cavities in each structure. Ligands, ions, and water molecules were first removed from the corresponding mmCIF file with MDAnalysis [44]. The system was then processed by mkgridXf with inner and outer radii spheres equal to 1 Å and 8 Å, respectively. The cavities found by mkgridXf were extremely large, often extending throughout the entire RNA structure. We therefore decided to further classify them into sub-cavities.
2. Sub-cavity classification. We segmented each cavity found by mkgridXf using watershed, an image processing algorithm that can be used to detect contours and separate adjacent objects in a 3D volume [50]. The minimum distance between the centers of two sub-cavities was set at 6 Å based on the dimension of the smallest ligands in our database.
3. Identification of sub-cavities occupied by ligands and volume calculation. We used Delaunay triangulation [51] to identify all the sub-cavities occupied by each ligand. To avoid including sub-cavities populated by a few ligand atoms, we considered a sub-cavity to be occupied only when at least 20% of the ligand atoms were found inside its volume. Finally, for each ligand we merged all the occupied sub-cavities and calculated the total volume. The use of different cutoffs for the sub-cavity occupation in the range 10%-30% did not significantly alter the final volume distribution (**Fig. S1**).

#### Pocket analysis with Schrodinger suite

We analyzed each pocket in the HARIBOSS database using the tools available in Schrodinger Maestro suite v. 2021-3. This analysis is articulated in 4 steps:

1. We extracted the region within 12 Å of the ligand in order to optimize the computational cost of the analysis, as our database contained large systems up to 10^4^ kDa;
2. We used Maestro [52] to prepare each substructure by adding missing hydrogens, optimizing their assignment at pH of 7.4 with PROPKA [53], and minimizing the resulting model using steepest descent in combination with OPLS3 [54];
3. We used SiteMap [55, 56] to first define pockets using the coordinates of the bound ligand and a sitebox equal to 6 Å. In cases in which multiple pockets were identified for the same ligand, we used Delaunay triangulation to select the pockets occupied by at least 20% of the ligand atoms (**Table S2**).
4. For the pockets occupied by ligands, we analyzed: volume, number of site points, hydrophobicity, hydrophilicity, enclosure (or buriedness), donor/acceptor character, SiteScore, and Dscore. Except for the volume, these are unitless quantities whose ranges were calibrated on a benchmark set of protein-ligand complexes. In particular, SiteScore and Dscore quantify the propensity of a pocket to bind ligands and drug-like molecules, respectively. Both scoring functions are defined in terms of pocket size (*n*), enclosure (*e*), and hydrophilicity (*p*) as:

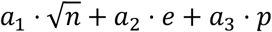

with different coefficients for SiteScore (*a*_1_= 0.0733; *a*_2_ = 0.6688; *a*_3_= −0.20) and Dscore (*a*1 = 0.094; *a*2 = 0.60 ; *a*3 = 0.324).

### HARIBOSS website

The HARIBOSS database is available online through a web application (https://hariboss.pasteur.cloud). The online version of HARIBOSS is enriched with additional cross-links and properties, either fetched from RCSB PDB or computed using the RDKit library^1^. The web application allows to query, visualize and download the data using either a compound-centric or a complex-centric perspective.

The compound-centric perspective (https://hariboss.pasteur.cloud/compounds/) allows to access the list of compounds identified as RNA binders in HARIBOSS. Multiple options allow to filter these compounds based on their properties (e.g. molecular weight) or on the properties of the PDB complexes where they have been identified (e.g. experimental resolution). It is possible to visualize the compounds either as thumbnails (the default representation), as a list or as a table. The details of each of these compounds include different identifiers (SMILES, IUPAC, InChi, InChiKey), as well as links to external databases, compliance with some of the drug-likeness criteria, and the list of RNA-SM complexes in which each compound has been identified. Multiple elements of this section of the web application were heavily inspired by the iPPI-DB database [57]. The graphical representation of the compounds uses the SmilesDrawer component [58].

The complex-centric perspective (https://hariboss.pasteur.cloud/complexes/) provides an access to the list of RNA-SM complexes. It also includes filtering options based on the properties of the complexes, as well as of the associated ligands. For each complex, a detail page provides a graphical representation of the complex, using the NGL library [59], and its main properties. This page also lists the identified pockets, color-coded according to their SiteScore ligandability score. Selecting the pockets allows to highlight them in the graphical representation of the complex. Both the compound- and complex-centric perspectives provide similar download features to retrieve either the entire query represented on the screen or a set of items that can be selected using the corresponding checkboxes.

## Results and discussions

HARIBOSS is constructed in 3 steps (**Fig. 1A**, Materials and Methods): initial fetching of RNA structures from the PDB, filtering of the database to identify systems with at least one small molecule bound to RNA (redundant HARIBOSS), and clustering based on RNA sequence identity and pocket structural similarity (non-redundant HARIBOSS). As of May 2022, the redundant version of HARIBOSS contains 716 PDB structures of RNA-SM complexes, for a total of 1226 pockets occupied by 267 unique ligands. After clustering, the non-redundant version of HARIBOSS contains 484 PDB structures, with a total of 676 pockets occupied by 267 unique ligands. In the following paragraphs, we present an overview of the general properties of the structures included in the HARIBOSS database, the physico-chemical properties of the ligands bound to RNA, and of the pockets and cavities.

### General properties of the HARIBOSS structures

The molecular composition of the structures changed at different stages of the construction of the database. Initially, HARIBOSS contains systems composed of RNA/protein complexes (59.3%), RNA molecules only (23.1%), RNA/protein/DNA (13.4%) and RNA/DNA complexes (4.2%) (**Fig. 1A, cyan**). All the RNA/protein/DNA complexes along with several RNA/protein complexes were filtered out because molecules other than RNA were involved in ligand binding and therefore were beyond the scope of the present study (**Fig. 1A, red**). In the non-redundant version of HARIBOSS, the majority of structures (62.0%) contained only RNA molecules, while the remaining part consisted of either RNA/protein (37.8%) or RNA/DNA (0.2%) complexes. (**Fig. 1A, yellow**). The systems initially included in our database significantly vary in size, with a total mass ranging from 0.5 to 10^4^ kDa regardless of the presence of a small molecule (**Fig. 1B, cyan**). Interestingly, all the structures above 10^2^ kDa are ribosomal RNA/protein complexes. Furthermore, the filtering of the initial database reduced the number of structures with mass below 5 kDa from 23% to 9% (**Fig. 1B, red**). This significant reduction supports the idea that RNA must have sufficient structural complexity to bind small molecules [14, 18, 40].

The majority of structures in the non-redundant version of HARIBOSS (∼83%) were determined by X-ray crystallography with a typical resolution of 2.8 Å. However, over the past 5 years the number of cryo-electron microscopy (cryo-EM) structures, in particular of large ribosomal RNA/protein complexes, has steadily been increasing (**Fig. 1C**) and currently represents 11% of our database with a typical resolution of 3.0 Å. Finally, structures determined by Nuclear Magnetic Resonance (NMR) spectroscopy constitutes the remaining 6% of the database. Cryo-EM and NMR structures are particularly interesting from the point of view of rational drug design as they provide dynamic information about the RNA molecules that can be exploited in virtual screenings [60]. Overall, the number of RNA-SM structures deposited in the PDB every year is constantly increasing as a reflection of the growing interest of the community in studying RNA-SM interactions (**Fig. 1C**).

### Properties of the HARTBOSS ligands

To explore the chemical space of RNA binders, we first created a dataset of unique ligands, based on their PDB identifier. As of May 2022, the total number of unique ligands in the HARIBOSS database is 267. Among these ligands, there are 11 that appear in more than 10 structures of the non-redundant HARIBOSS (**Table S3**). To provide an overview of the chemical matter present in our dataset, we used a minimum spanning tree representation (Materials and Methods) (**Fig. 2A**). Diverse ligand scaffolds are present, with some of them belonging to known classes of therapeutic agents. There is a significant number of structures of known antibiotics, like Linezolid and other oxazolidinones, Tiamulin, Eravacycline and other tetracyclines (**Fig. S2**). It is not surprising that GTP and nucleoside analogs as well as long polar molecules, like PEG and spermidine-derived polyamines, also appeared as common RNA binders.

**Figure 2.**
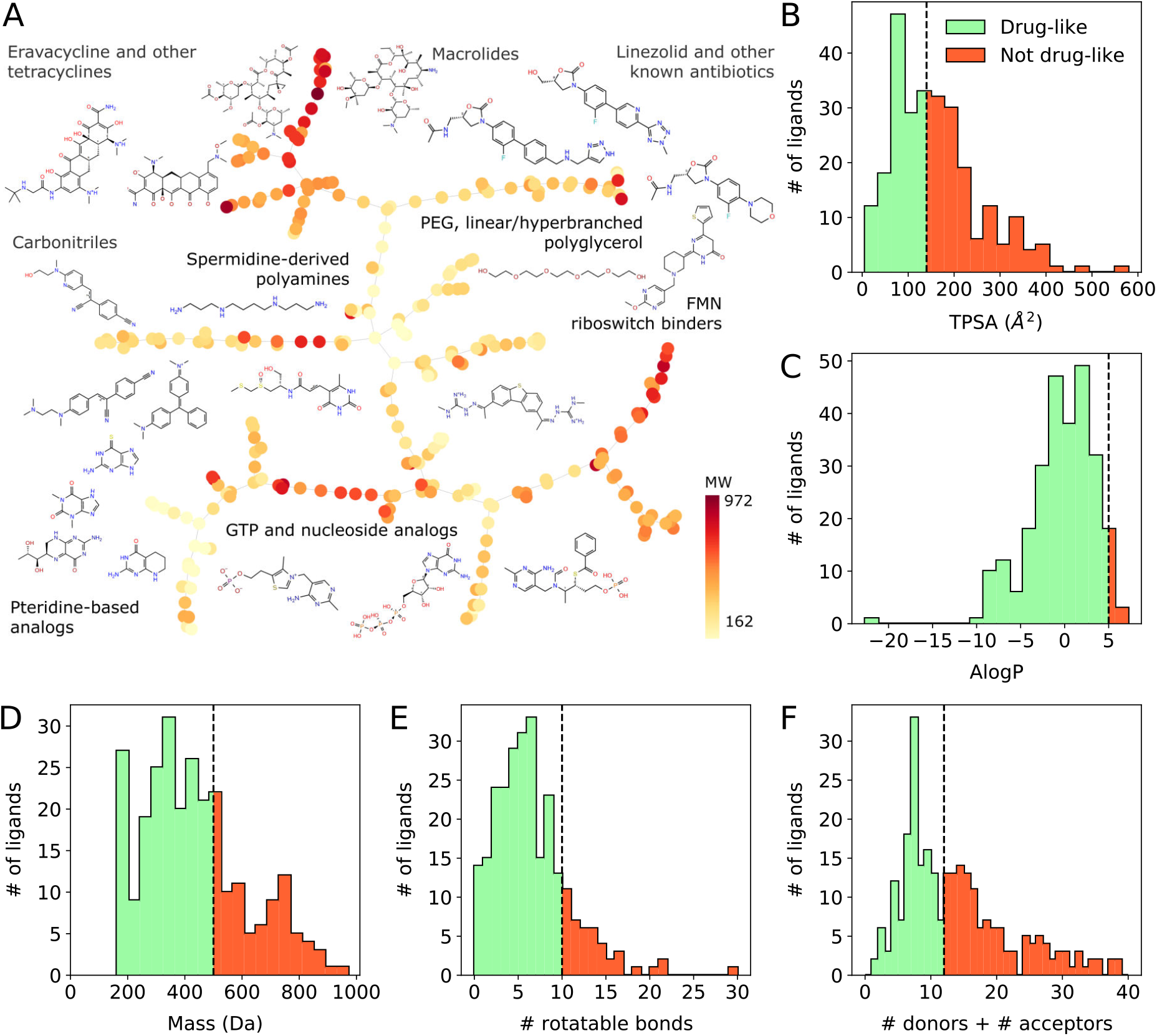
Pharmaco-chemical properties of the ligands in the HARIBOSS entries. A) 2-D representation of the chemical space of the 267 unique RNA binders obtained by an undirected weighted c-approximate k-nearest neighbor graph (Materials and Methods). Each node represents a ligand and is colored by its molecular weight (MW). B - F) Distributions of pharmaco-chemical properties of the ligands: the topological polar surface area (TPSA, B), the octanol-water partition coefficient (AlogP, C), the molecular mass (D), the number of rotatable bonds (E), and the sum of hydrogen bond donors and acceptors (F). For each panel, the dashed line indicates the threshold that defines drug-like compounds based on Lipinski’s rule of 5 f61l or Veber rule f62l. Green/red portions of the histogram represent the regions satisfying/violating these criteria.

The high polarity of RNA binders is depicted in their TPSA distribution, with less than half of the molecules (44%) being beyond the Veber drug-likeness threshold (140 Å^2^) (**Fig. 2B**). This high polarity is in line with the overall good solubility of the ligands in our HARIBOSS dataset (**Fig. 2C**). The compounds in our database significantly vary in size with a molecular mass spanning a range from 162 Da to 972 Da. The vast majority of these compounds (82%) have a mass lower than 600 Da (**Fig. 2D**). Approximately 30% of the RNA binders had a number of either donors and/or acceptors above the drug-likeness threshold (**Fig. 2E** and **Fig. S3**). This is again consistent with the high polar nature of these small molecules as well as the RNA targets. Moreover, RNA structurally forms “warm-like” long cavities and therefore long and flexible small molecules are expected to be among its binders. This is confirmed by the distribution of the total number of rotatable bonds of the HARIBOSS small molecules, which in 12.4% of the cases exceed the Lipinski’s threshold of 10 rotatable bonds (**Fig. 2F**).

All the above-mentioned properties are reported also on the HARIBOSS website (https://hariboss.pasteur.cloud). To complement the analysis of the small-molecule chemical space, QikProp (Schrordinger Suite 2021.v3) was used to calculate the following properties: solvent-accessible surface area (SASA) and its hydrophobic and hydrophilic content (FOSA and FISA respectively), predicted IC50 value for blockage of hERG K+ channels (QPlogHERG), Caco-2 cell permeability (QPPCaco) and brain/blood partition coefficient (QPlogBB). Based on this analysis, SASA was found, as expected, to be proportional to the MW with the exception of very large molecules (MW>600) in which the ligand conformation may significantly alter SASA (**Fig. S4**). On the contrary, weaker correlations were found between MW and both FOSA (**Fig. S5**) and FISA (**Fig. S6**), and between FISA and FOSA (**Fig. S7**). Interestingly, 25.2% of our ligand database is above the FISA drug-like threshold, while only 3% shows high FOSA, in agreement with TPSA (**Fig. 2B**). Among the set of unique RNA binders 55% do not show potential hERG liabilities (**Fig. S8**). Approximately half of the ligands (52%) are predicted to have poor Caco-2 cell permeability (**Fig. S9**) while more 70% of the molecules have a good predicted brain/blood partition coefficient (**Fig. S10**).

### Properties of the RNA pockets and cavities

In the non-redundant version of HARIBOSS, the majority of pockets occupied by ligands and identified by SiteMap and mkgridXf (Materials and Methods) are formed by a single RNA chain (67%), while the remaining are at the interface of two (29%) or more (4%) RNA chains (**Fig. 3A**, yellow bars). Overall, most of these pockets (67%) were considered to be potential ligand binding sites (ligandable) according to the SiteScore scoring function (SiteScore ≥ 0.8, **Fig. 3B**). However, according to the Dscore score, only 35% of the pockets were classified as druggable (Dscore ≥ 0.98), 24% as difficult targets (0.83 ≤ Dscore < 0.98), and the remaining 41% as undruggable (**Fig. 3C**). SiteScore and Dscore depend on several physico-chemical properties of the pocket and have been optimized for protein molecules. Among these properties, RNA pockets presented a typical hydrogen-bond donor/acceptor character (**Fig. 3D**) and hydrophilicity (**Fig. 3E**) similar to proteins.

**Figure 3.**
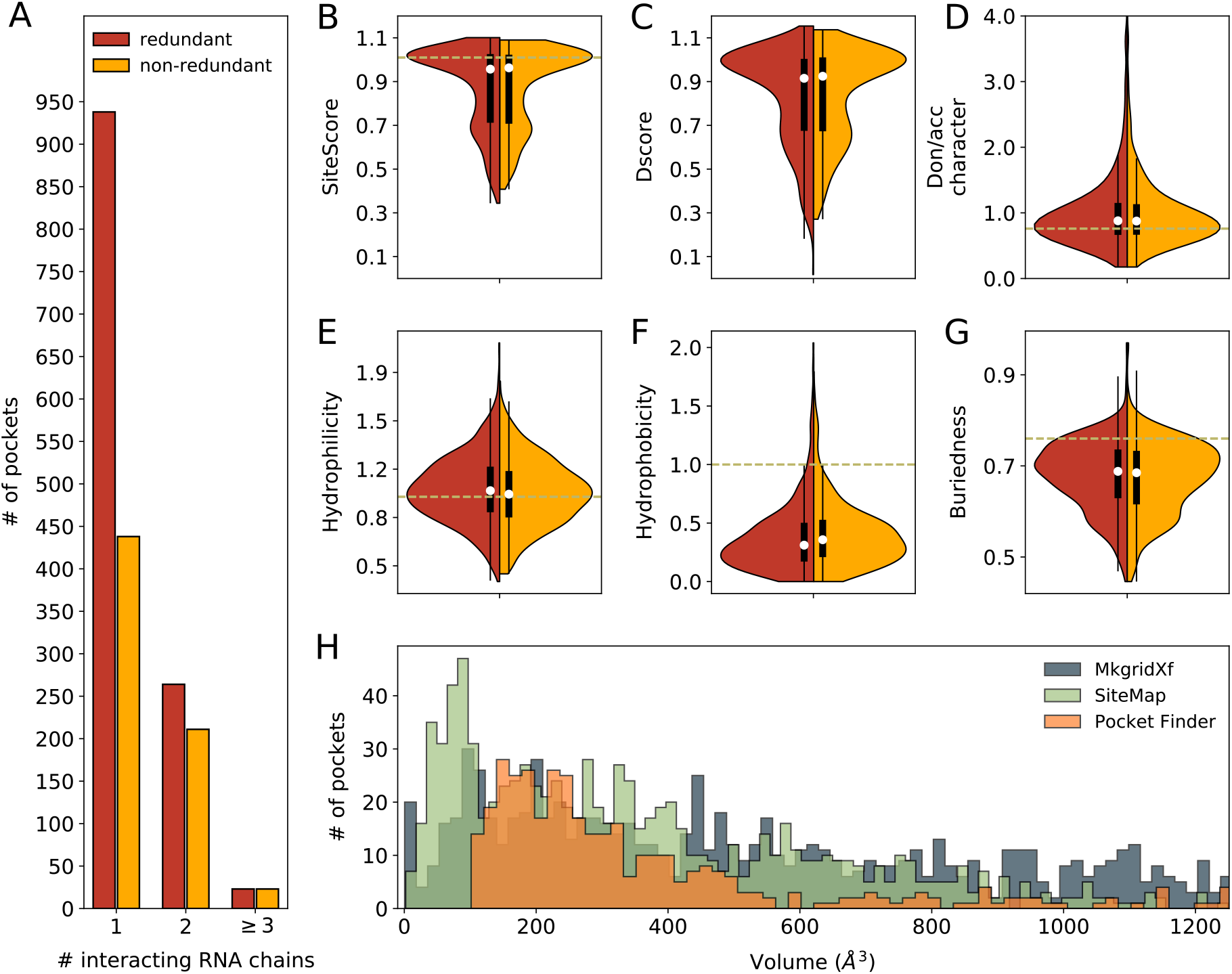
Physico-chemical properties of RNA-SM pockets. A) Number of RNA-SM pockets as a function of the number of interacting RNA chains in the redundant (red) and non-redundant (yellow) HARIBOSS database. B - G) Violin plots representing the distributions of properties of RNA-SM pockets calculated by SiteMap: ligandability (SiteScore, B) and druggability scores (Dscore, C), hydrogen bond donor/acceptor character (D), hydrophilicity (E), hydrophobicity (F), and buriedness (G). The average values of these properties calculated on a benchmark of protein-SM complexes is represented by dashed lines f55, 56l. H) Volume distribution of RNA-SM pockets calculated with SiteMap (light green) and mkgridXf [49] (dark green) on the redundant HARIBOSS database, and with PocketFinder (orange) in Hewitt *et al*. f40l. The x-axis is limited at 1250 A^3^.

However, RNA pockets appeared to be less hydrophobic (**Fig. 3F)** and more exposed to solvent (**Fig. 3G**). This different character is a consequence of the more polar nature of RNA molecules.

A significant number of pockets was considered non ligandable (SiteScore < 0.8, **Fig. 3B**). In many cases, low ligandability and druggability can be interpreted in terms of the physico-chemical properties of the pocket. For example, hydrophilic pockets particularly exposed to solvent were generally considered not druggable (**Fig. S11**), as in the case of the structure of a RNA primer3template bound to ligand 5GP (PDB 5dhb, **Fig. 4A**). In contrast, the hydrophobicity of RNA-SM pockets appears not to be strongly correlated with ligandability (**Fig. S12**). Highly hydrophobic pockets, like the site in which ligand EKM binds the Mango-II Fluorescent RNA Aptamer structure (PDB 6c64, **Fig. 4B**), and less hydrophobic pockets, such as the binding site of ligand UG4 in *Fusibacterium ulcerans* ZTP riboswitch (PDB 6wzs, **Fig. 4C**), were found to be equally ligandable. However, druggable pockets have a tendency to be more hydrophobic (**Fig. S13**).

**Figure 4.**
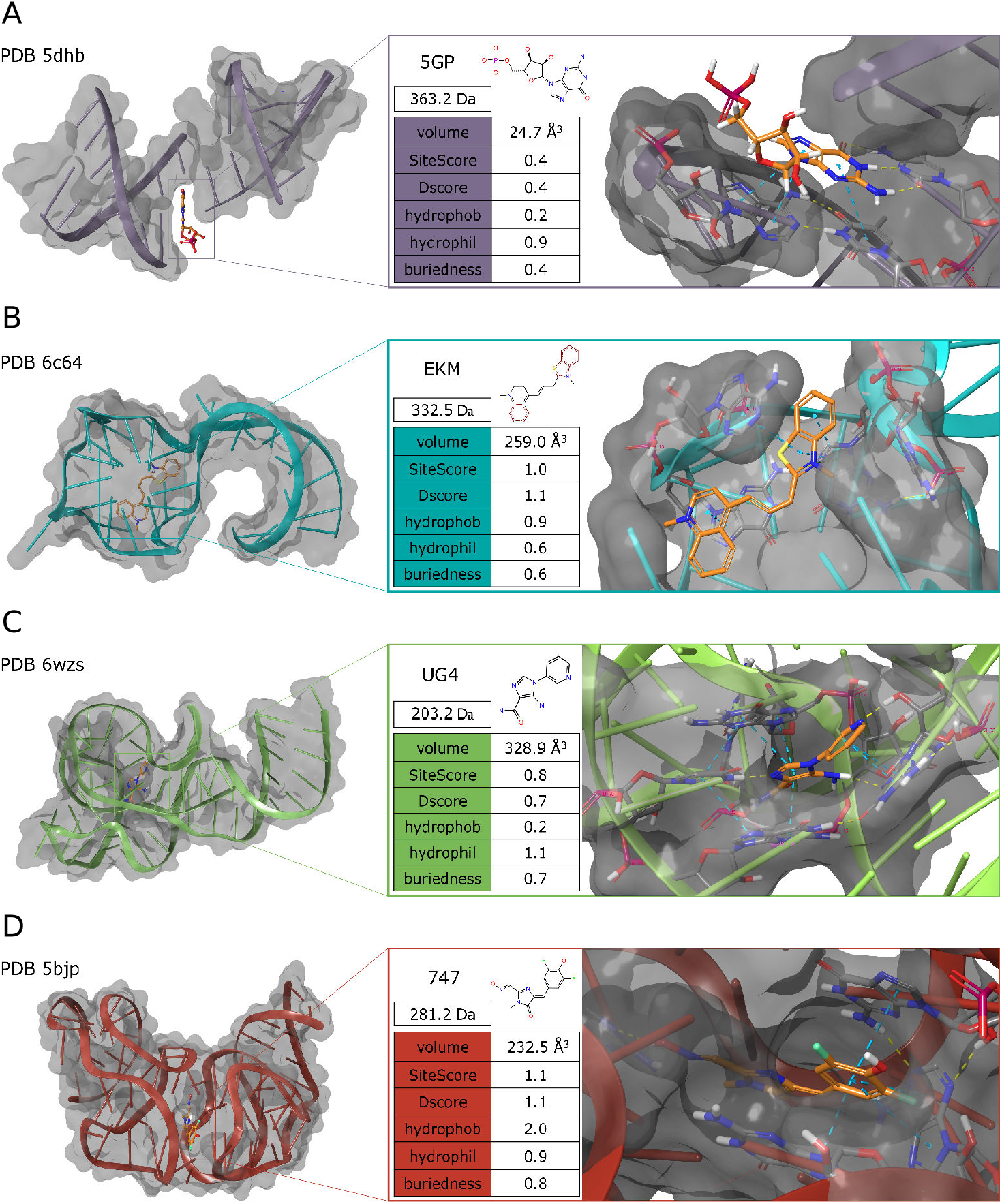
Examples of RNA-SM structures included in HARIBOSS: a RNA primer-template bound to ligand 5GP (PDB 5dhb, A), the Mango-II Fluorescent RNA aptamer bound to ligand EKM (PDB 6c64, B), the *Fusibacterium ulcerans* ZTP riboswitch bound to UG4 (PDB 6wzs, C), and the Corn RNA aptamer bound to ligand 747 (PDB 5bj, D). Left column: PDB code and structure of the RNA-SM complex. Right column: name, molecular weight, 2D representation of the small molecule, table with ligand properties calculated by SiteMap, and close view of the pocket occupied by the small molecule. The following types of RNA-SM interactions are highlighted by dashed lines: hydrogen bonds (yellow), salt bridges (purple), and π-π stacking (cyan).

In agreement with previous studies [40], RNA-SM pockets span a broad range of volumes, independently from the software and dataset used for pocket identification and volume calculation (**Fig. 3H**). The size of the most druggable and ligandable pocket in HARIBOSS, corresponding to the binding site of 747 to the Corn RNA aptamer (PDB 5bjp and 5bjo, **Fig. 4D**), is equal to ∼230 Å^3^. However, cavities larger than 300 Å^3^ were also classified as ligandable (**Fig. S14**). Given the size of the small molecules in HARIBOSS (**Fig. 2A**), these large cavities cannot be fully occupied by a ligand. Indeed, although for small pockets the ligand volume is often bigger than the pocket volume, suggesting that the ligand is exposed to water, for larger pockets the ligand volume is consistently smaller compared to the pocket volume (**Fig. S15**). Further investigations are needed to understand whether this finding is due to the fact that RNA molecules form large cavities that are only partly occupied by ligands or by artifacts of the software used for detecting cavities and calculate their volume. Finally, all cavities with volume smaller than 100 Å^3^ were considered neither druggable (**Fig. S14**) nor ligandable (**Fig. S16**).

## Conclusions

Here, we presented HARIBOSS, a curated database of structures of RNA-small molecule complexes, built to aid the development of computational drug design pipelines. For each HARIBOSS entry, we provided general structural information and we analyzed the physico-chemical properties of the ligands bound to RNA, and of the respective pockets. For the majority of structures in our database, the experimentally-resolved pockets were confirmed as ligandable. Only one third of all pockets was ranked as good for drug design purposes, the remainder part being a difficult target or undruggable. Our analysis indicates that low druggability is due to the fact that RNA pockets are less hydrophobic and more exposed to solvent than protein pockets. In line with these findings, RNA binders in the HARIBOSS database were mostly highly polar and water-soluble ligands. Known classes of antibiotics and endogenous polar ligands are among the most frequent RNA binders in PDB. Cell permeability is, as expected, a major issue for a significant part of these molecules. Future studies will be aimed at identifying HARIBOSS compounds that are RNA-selective and characterizing the physico-chemical interactions that determine their selectivity.

As of today, HARIBOSS contains only static RNA-SM structures. This is already an important step that will aid the discovery of RNA binders using 3D structure-based, rational drug discovery approaches. However, this is only the first step in the identification of RNA inhibitors and modulators. Demonstrating that binding translates to changes in dynamics and function is the most challenging part. The next natural step is to include in HARIBOSS structural ensembles representing the highly-dynamic nature of RNA. Characterizing the dynamic properties of RNA molecules is particularly important to identify selective and specific compounds using structure-based approaches that target individual members of RNA conformational ensembles [60].

HARIBOSS can be accessed via a web interface available at https://hariboss.pasteur.cloud and explored using a compound or pocket perspective. Our database will facilitate: *i)* assessing the accuracy of existing protein-oriented drug design computational tools, identifying areas of improvement, and optimizing them for RNA molecules; *ii)* investigating the nature of RNA-SM interactions; *iii)* defining the chemical space of RNA binders and their potential to be used as drugs; *iv)* identifying new potential RNA targets based on pocket druggability or starting from a specific compound. In conclusion, our comprehensive, curated, and regularly updated database of RNA-SM structures is a stepping stone for the scientific community to develop novel *in silico* approaches to discover compounds for direct RNA targeting.

## Acknowledgments

We would like to thank Arnaud Blondel and Guillaume Bouvier (Institut Pasteur, Paris) for assistance with mkgridXf and watershed calculations. The development of the HARIBOSS website built on the web infrastructure developed for the iPPI-DB database, a project led by Olivier Sperandio (Institut Pasteur, Paris). This work used the Kubernetes cluster provided by the IT department at Institut Pasteur, Paris. We thank Thomas Ménard for his help with the Kubernetes configuration, Fabien Mareuil for his help with the use of the NGL javascript library.

## Footnotes

1 https://doi.org/10.5281/zenodo.591637

