## Supplementary Figures and Tables for "HARIBOSS: a curated database of RNA-small molecules structures to aid rational drug design"

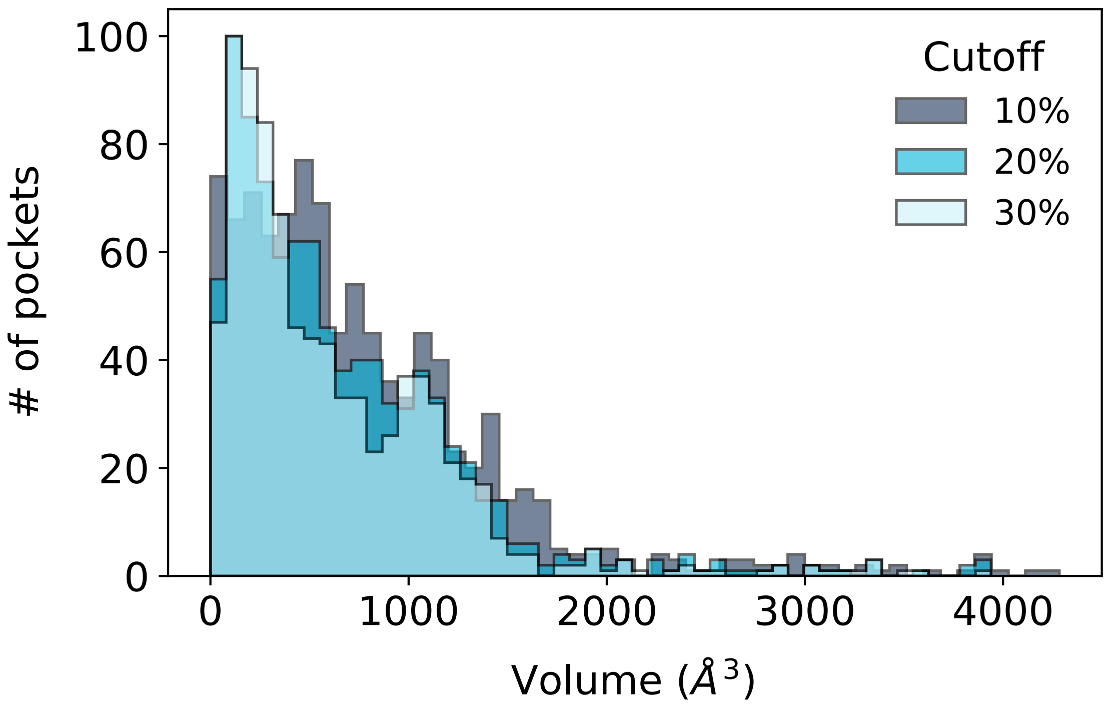


**Figure S1. Cavity analysis with mkgridXf.** Volume distributions of the cavities found by mkgridXf on the redundant HARIBOSS database using a cutoff for the sub-cavity occupation equal to 10% (dark blue), 20% (cyan), and 30% (light blue).

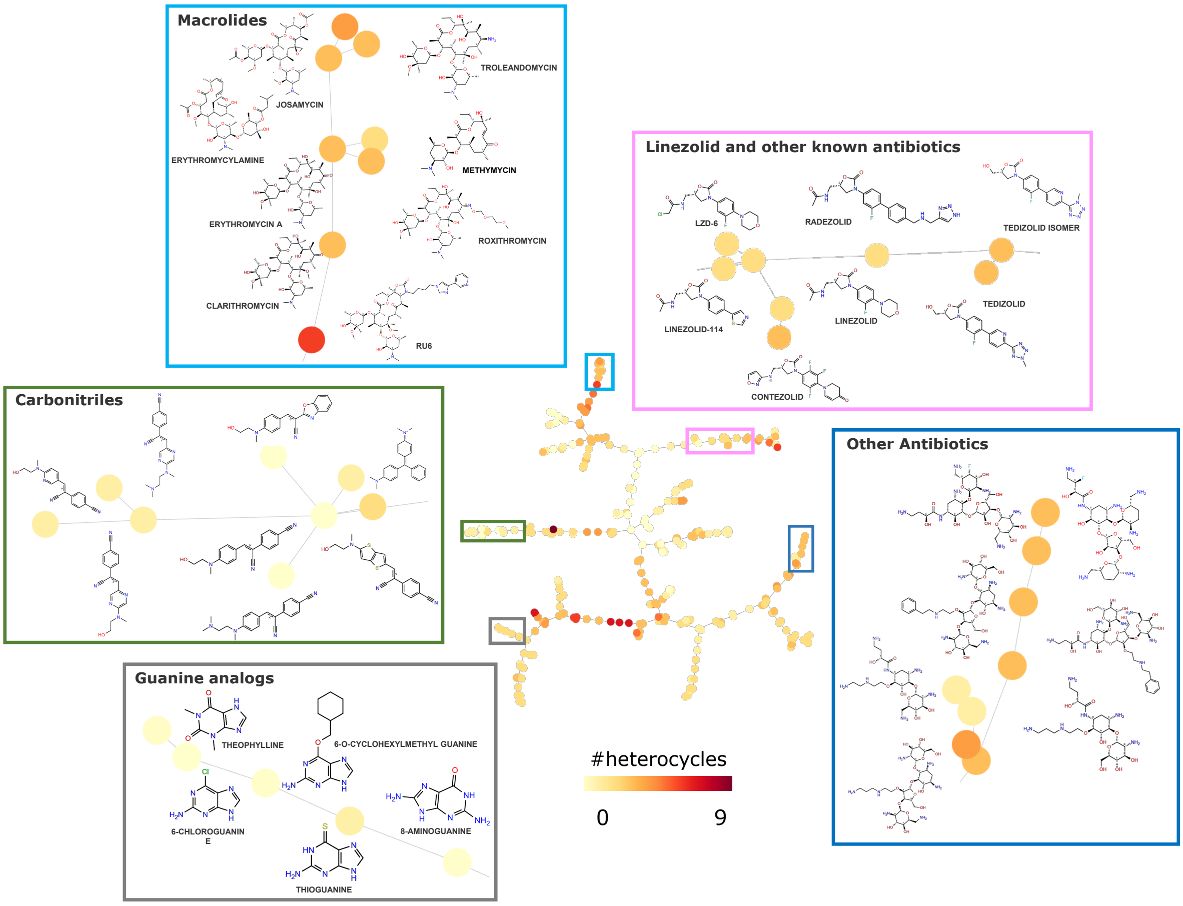


**Figure S2. Minimum spanning tree representation of the HARIBOSS small molecule database.** Five characteristic families of RNA binders belonging to the highlighted branches are shown in detail. Each node is colored according to the number of heterocycles present in the specific molecule.

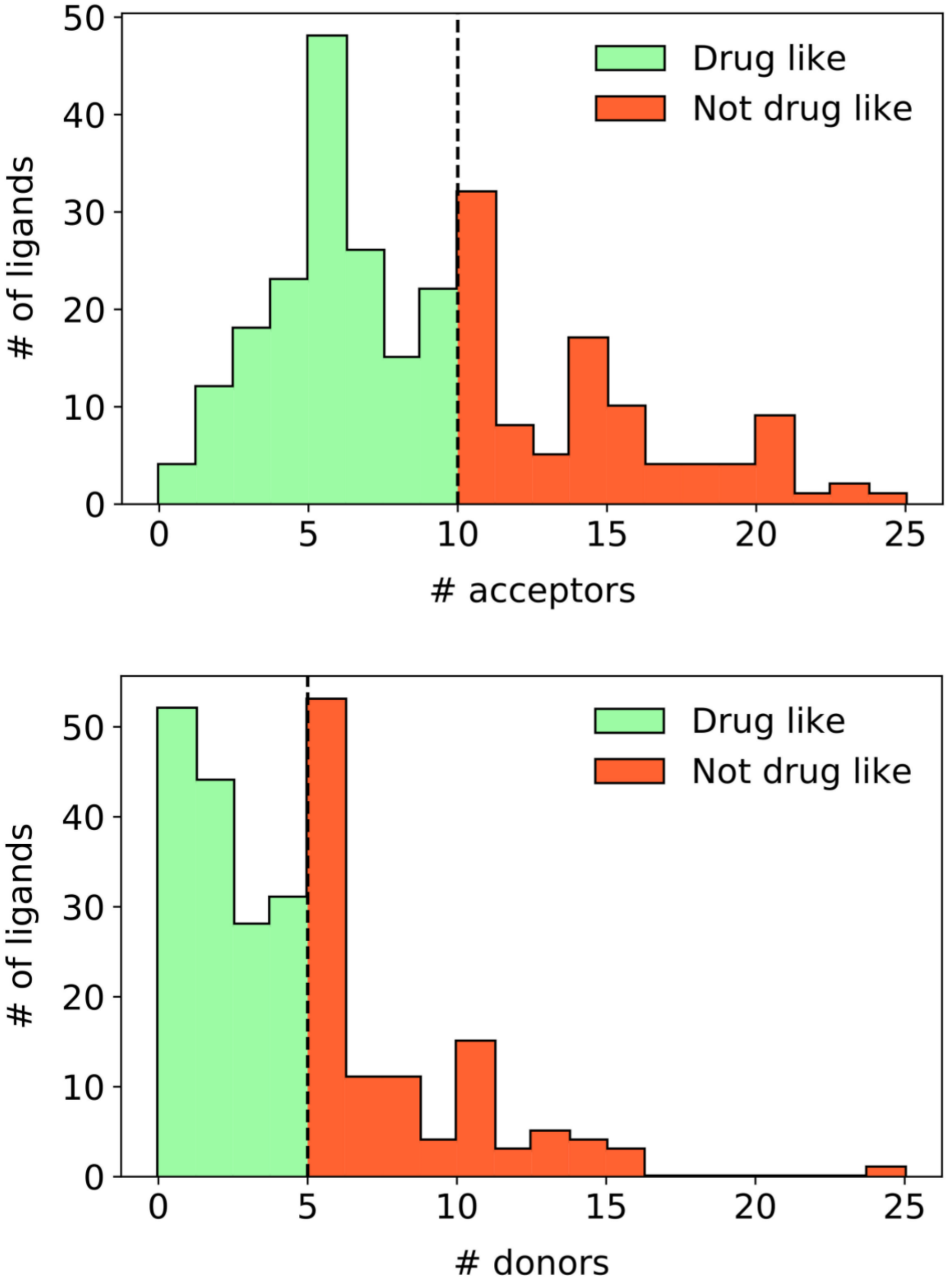


**Figure S3. Distribution of the number of hydrogen bond donors (top) and hydrogen bond acceptors (bottom) of the HARIBOSS ligands.** The dashed line indicates the threshold that defines drug-like compounds based on Lipinski's rule of 5 [1] or Veber rule [2]. Green/red portions of the histogram represent the regions satisfying/violating these criteria.

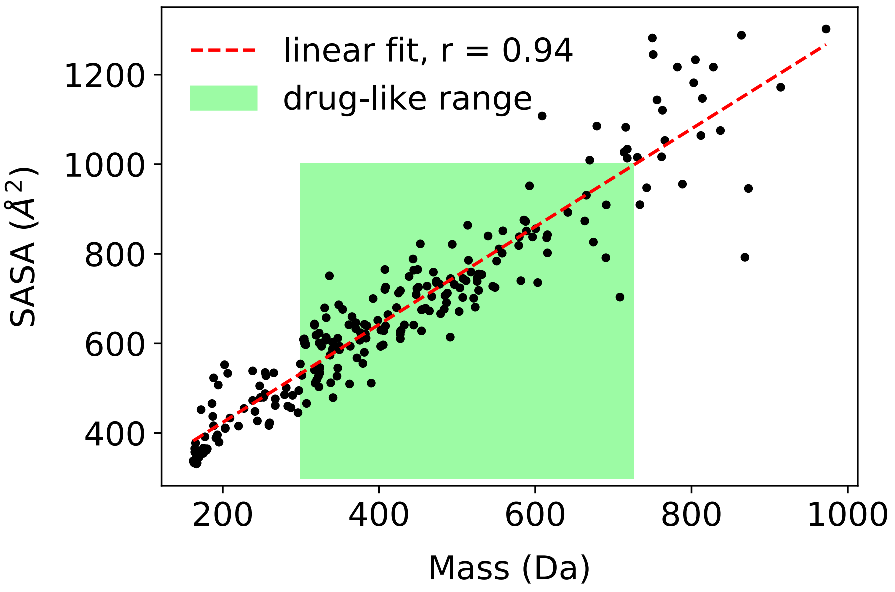


**Figure S4. Scatter plot of ligand mass vs Solvent Accessible Surface Area (SASA).** The properties were calculated on the set of unique ligands using QikProp. The green rectangle indicates the range of values corresponding to drug-like molecules as defined in [3], the red dotted line the linear fit.

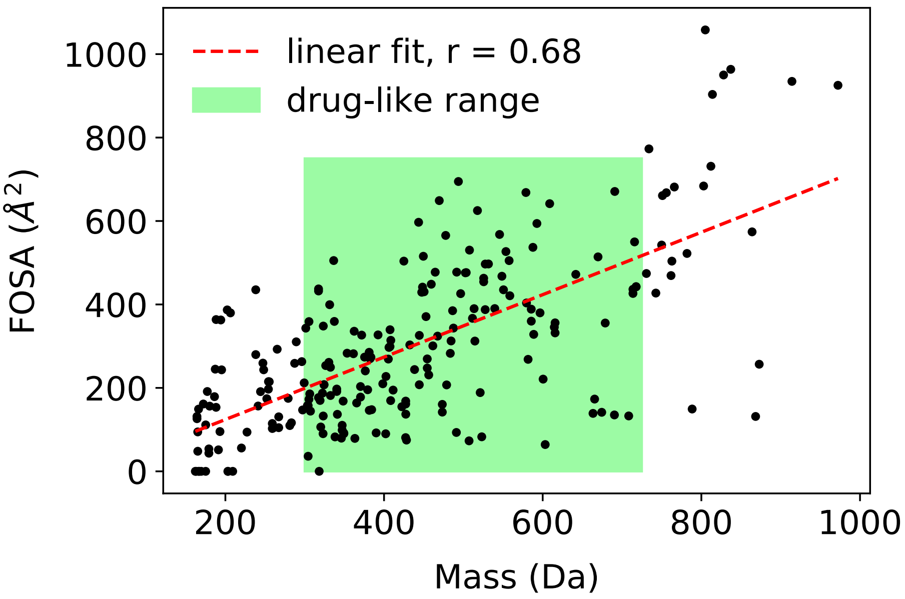


**Figure S5. Scatter plot of ligand mass vs Hydrophobic solvent accessible surface area (FOSA).** The properties were calculated on the set of unique ligands using QikProp. The green rectangle indicates the range of values corresponding to drug-like molecules as defined in [3], the red dotted line the linear fit.

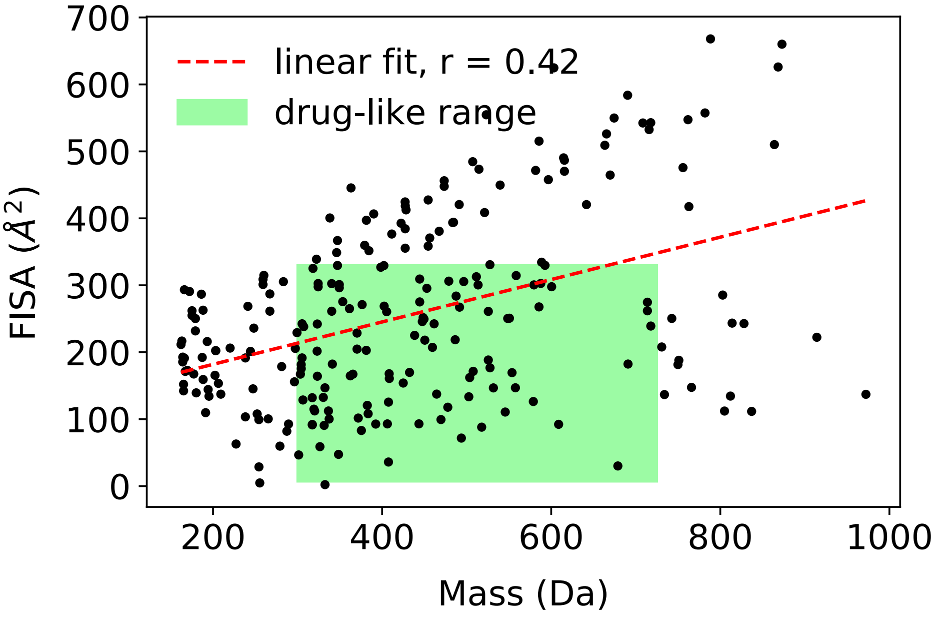


**Figure S6. Scatter plot of ligand mass vs Hydrophilic solvent accessible surface area (FISA).** The properties were calculated on the set of unique ligands using QikProp. The green rectangle indicates the range of values corresponding to drug-like molecules as defined in [3], the red dotted line the linear fit.

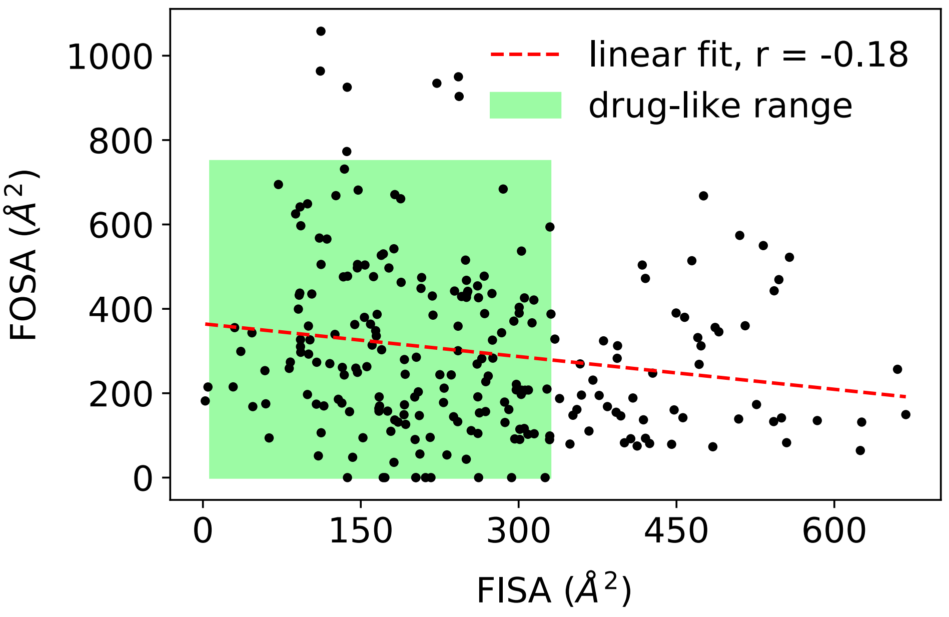


**Figure S7. Scatter plot of ligand FISA vs FOSA.** The properties were calculated on the set of unique ligands using QikProp. The green rectangle indicates the range of values corresponding to drug-like molecules as defined in [3], the red dotted line the linear fit.

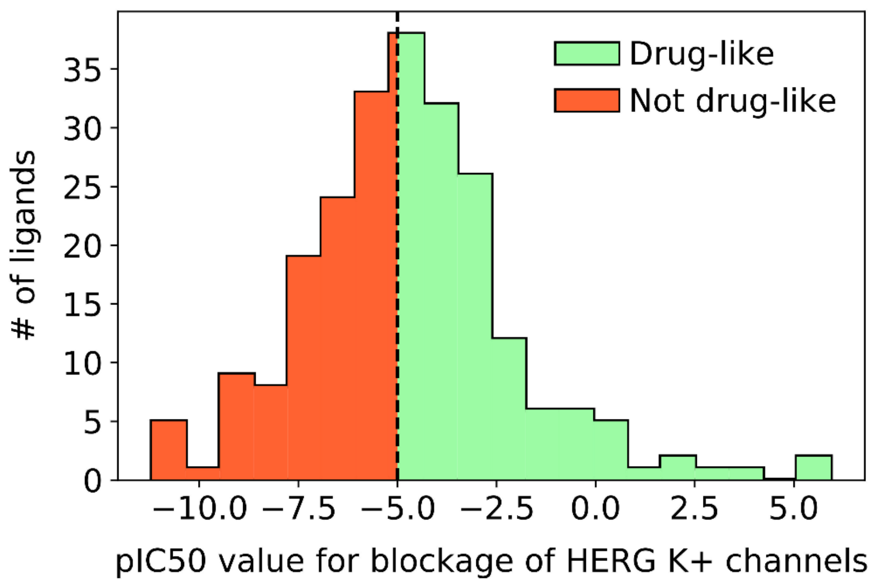


**Figure S8. Distribution of the predicted IC50 value for blockage of HERG K+ channels (QPlogHERG) of the HARIBOSS ligands.** The property was calculated on the set of unique ligands using QikProp. Green/red portions of the histogram represent the regions satisfying/violating the drug-likeness criterion for QPlogHERG as defined in [3].

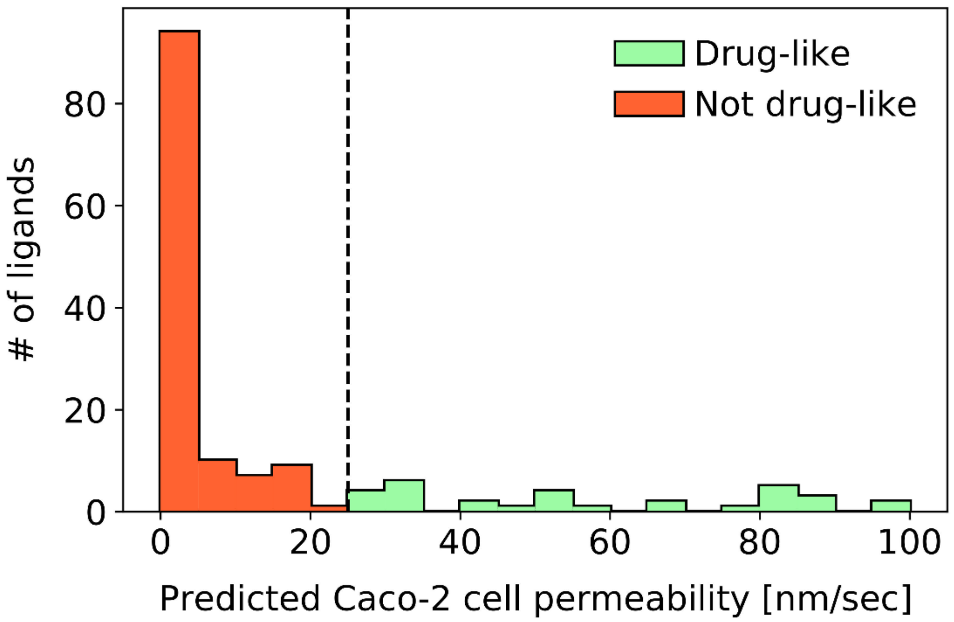


**Figure S9. Distribution of the predicted Caco-2 cell permeability (QPPCaco) of the HARIBOSS ligands.** The property was calculated on the set of unique ligands using QikProp. Green/red portions of the histogram represent the regions satisfying/violating the drug-likeness criterion for QPPCaco as defined in [3].

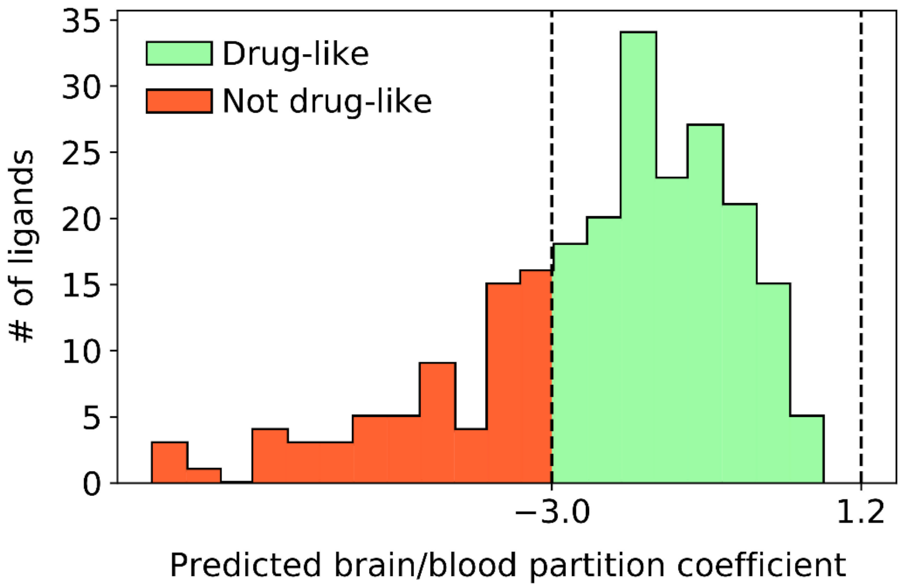


**Figure S10. Distribution of the predicted brain/blood partition coefficient (QPlogBB) of the HARIBOSS ligands.** The property was calculated on the set of unique ligands using QikProp. Green/red portions of the histogram represent the regions satisfying/violating the drug-likeness criterion for QPlogHERG as defined in [3].

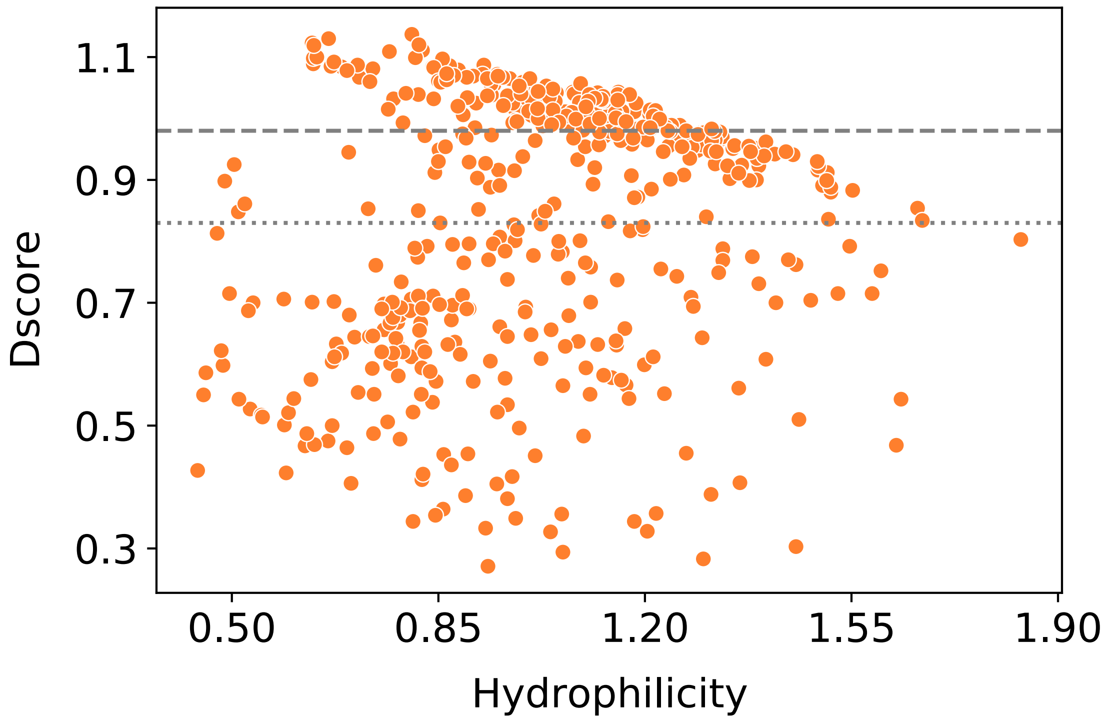


**Figure S11. Scatter plot of pocket hydrophilicity vs druggability score (Dscore).** The properties were calculated on the non-redundant HARIBOSS database using SiteMap. The dashed and dotted lines represent the thresholds for druggable and difficult-target pockets, respectively.

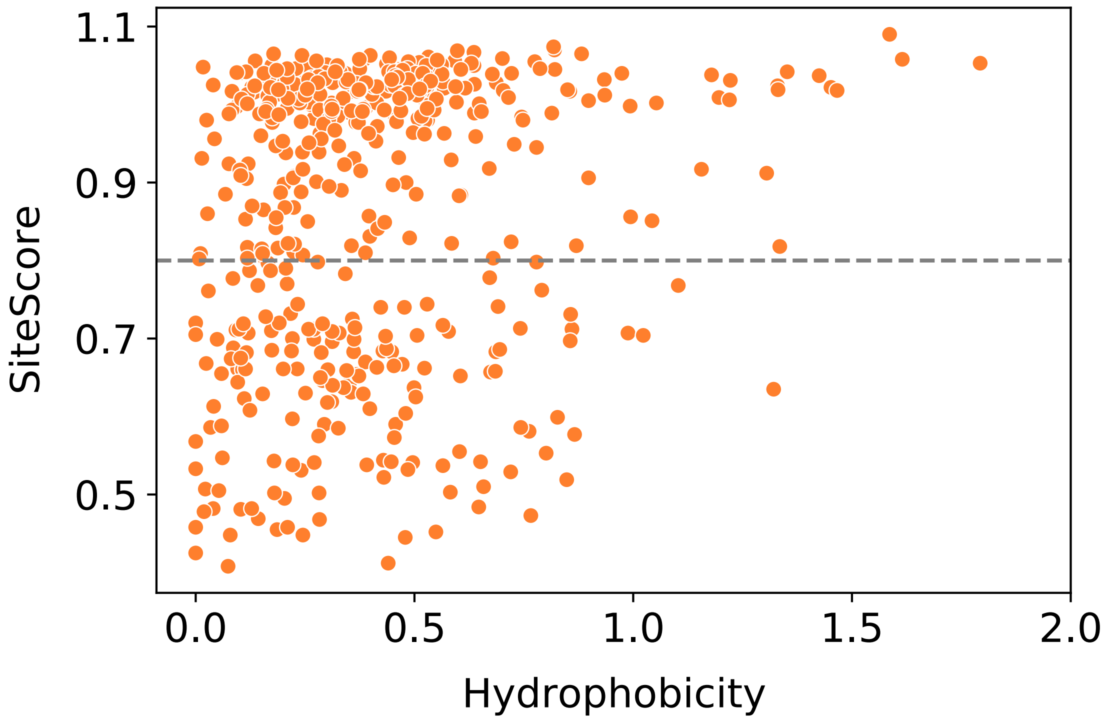


**Figure S12. Scatter plot of pocket hydrophobicity vs ligandability score (SiteScore).** The properties were calculated on the non-redundant HARIBOSS database using SiteMap. The dashed line represents the threshold for ligandable pocket.

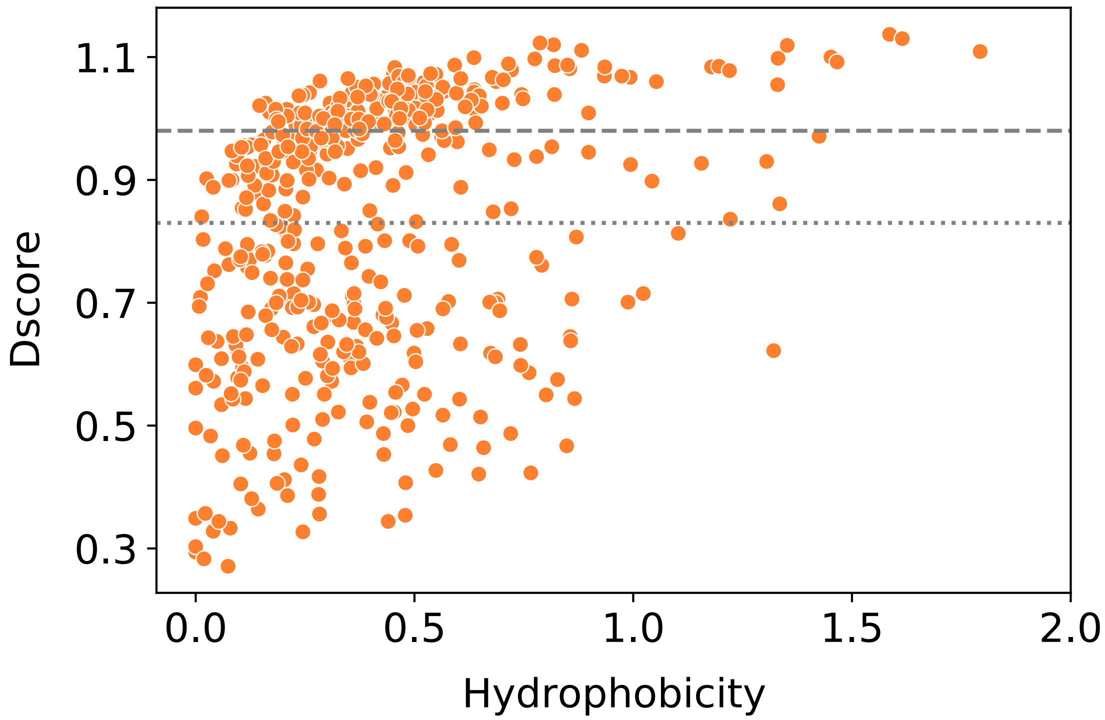


**Figure S13. Scatter plot of pocket hydrophobicity vs druggability score (Dscore).** The properties were calculated on the non-redundant HARIBOSS database using SiteMap. The dashed and dotted lines represent the thresholds for druggable and difficult-target pockets, respectively.

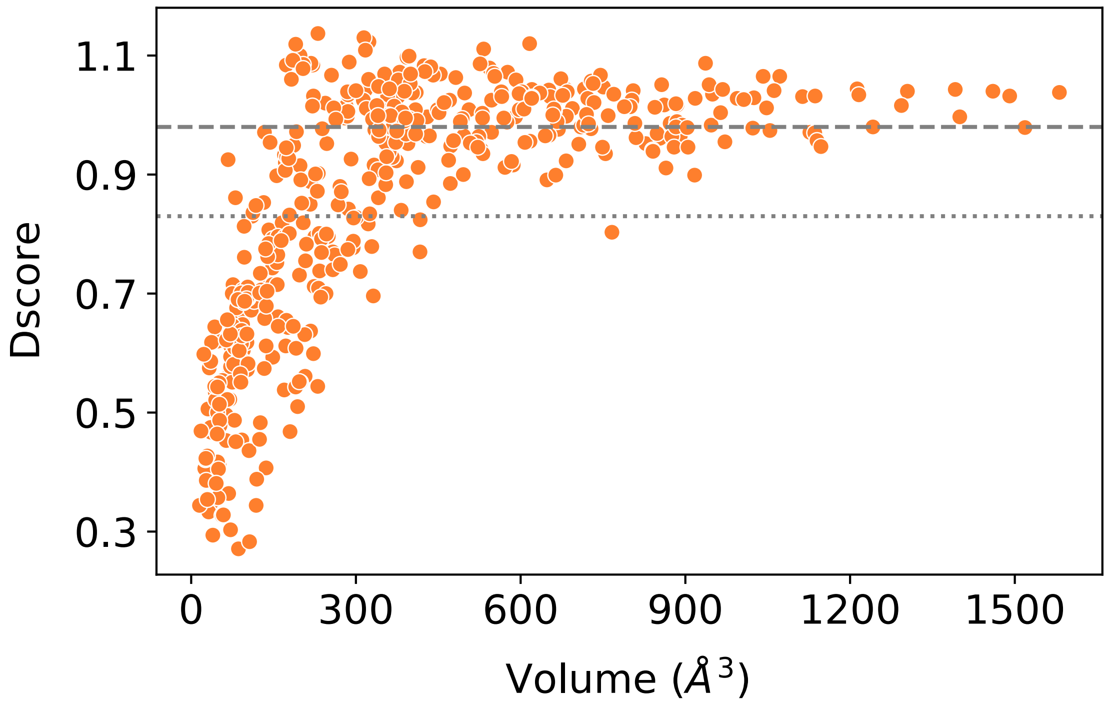


**Figure S14. Scatter plot of pocket volume vs druggability score (Dscore).** The properties were calculated on the non-redundant HARIBOSS database using SiteMap. The dashed and dotted lines represent the thresholds for druggable and difficult-target pockets, respectively.

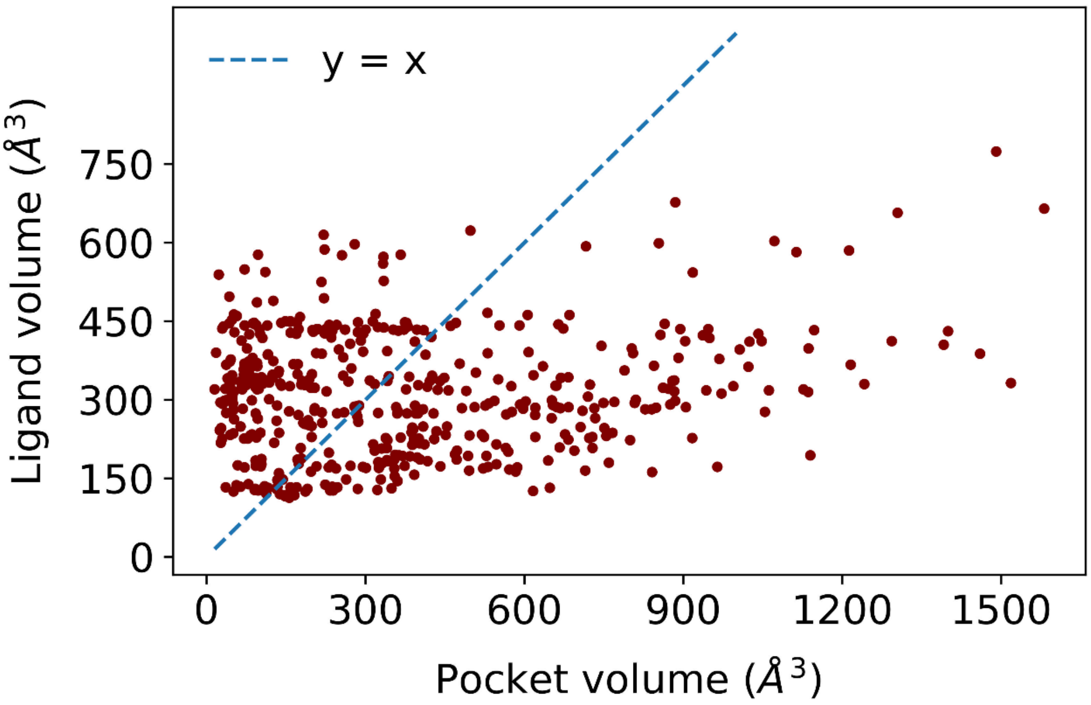


**Figure S15. Scatter plot of pocket volume vs ligand volume.** The properties were calculated on the non-redundant HARIBOSS database using SiteMap and the volume calculation script from Schrodinger Suite.

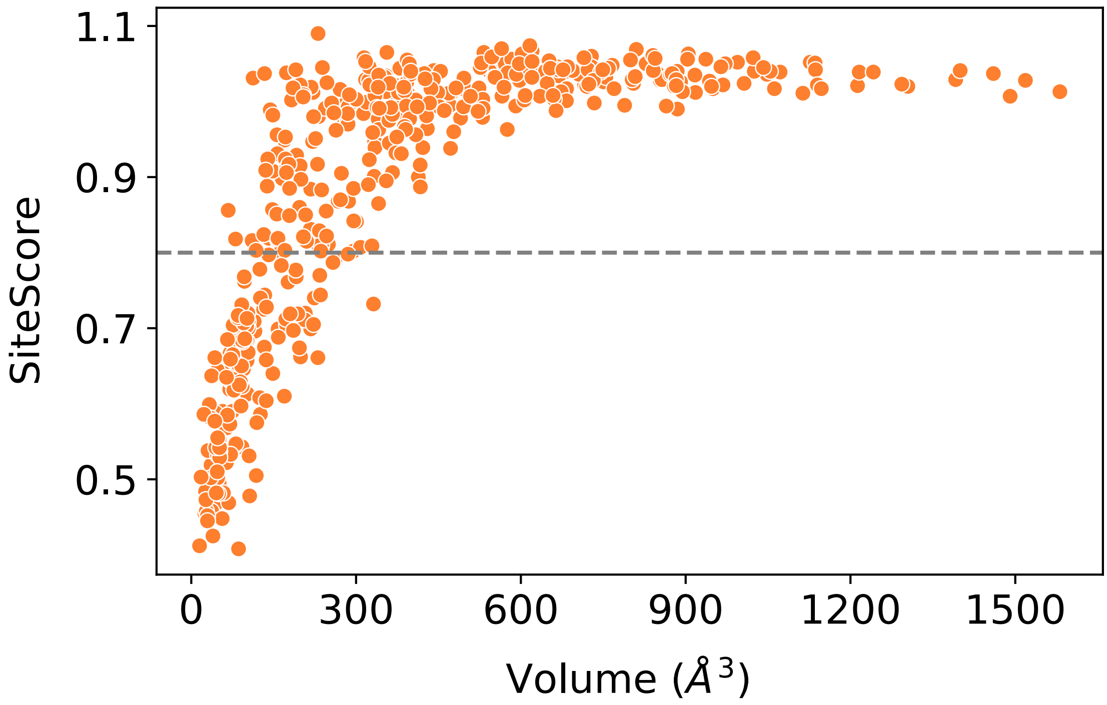


**Figure S16. Scatter plot of pocket volume vs ligandability score (SiteScore).** The properties were calculated on the non-redundant HARIBOSS database using SiteMap. The dashed line represents the threshold for ligandable pocket.

**Supplementary Tables**

| Cutoff | Redundant HARIBOSS # of interacting RNA chains | | | | | Non-redundant HARIBOSS # of interacting RNA chains | | | | |
|---|---|---|---|---|---|---|---|---|---|---|
| | **Total** | **1** | **2** | **3** | **4** | **Total** | **1** | **2** | **3** | **4** |
| **0** | 1158 | 794 | 323 | 38 | 3 | 610 | 380 | 211 | 17 | 2 |
| **5** | 1158 | 863 | 258 | 34 | 3 | 579 | 393 | 170 | 14 | 2 |
| **10** | 1145 | 873 | 248 | 23 | 1 | 573 | 405 | 159 | 9 | 0 |
| **20** | 1151 | 915 | 235 | 1 | 0 | 564 | 420 | 144 | 0 | 0 |

**Table S1. Number of pockets in the redundant and non-redundant HARIBOSS databases,** as a function of the number of RNA interacting chains and the minimum number of atoms (cutoff) for a chain to be considered as interacting. This analysis was performed on the HARIBOSS database updated in February 2022.

| Pocket analysis stage | # cases |
|---|---|
| **Input** | 1226 |
| **Preparation output** | 1180 |
| **Evaluation** | 1017 |

| Pocket composition and occupancy | # cases |
|---|---|
| **1 pocket** | **809** |
| **2 subpockets, with only 1 populated** | **116** |
| **2 subpockets, none populated** | **21** |
| **2 subpockets, both populated** | **7** |
| **More than 2 subpockets** | **12** |
| **No pockets** | **52** |

**Table S2. Statistics of the pocket analysis by SiteMap.**

| Non-redundant HARIBOSS | | |
|---|---|---|
| **Ligand PDB ID** | **Occurrence** | **Name** |
| PAR | 50 | Paromomycin |
| SPM | 30 | Spermine |
| NMY | 24 | Neomycin |
| LLL | 20 | Gentamicin C1A |
| GP3 | 19 | Diguanosine-5'-Triphosphate |
| SAM | 17 | S-Adenosylmethionine (SAMe) |
| 8UZ | 16 | TC007 |
| GET | 16 | Geneticin (G418) |
| GTP | 15 | Guanosine-5'-triphosphate |
| AM2 | 13 | Apramycin |
| NEG | 10 | Negamycin |

**Table S3. Occurrence of the 15 most frequent ligands in non-redudant HARIBOSS.**
